# Transcriptomic responses of Galápagos finches to avian pox virus infection

**DOI:** 10.1101/2021.10.15.464582

**Authors:** Sabrina M. McNew, Janaí Yepez, C. Diana Loyola, Catherine Andreadis, Birgit Fessl

## Abstract

Emerging pathogens can have devastating effects on naïve hosts, but disease outcomes often vary among hosts. Comparing the cellular response of different host species to infection can provide insight into mechanisms of host defense and the basis of host susceptibility to disease. Here, we used RNA-seq to characterize the transcriptomic response of Darwin’s finches to avian poxvirus, which is introduced to the Galápagos Islands. We tested whether gene expression differs between infected and uninfected birds, and whether transcriptomic differences were related either to known antiviral mechanisms and/or the co-option of the host cellular environment by the virus. We compared two species, the medium ground finch (*Geospiza fortis*) and the vegetarian finch (*Platyspiza crassirostris*), to determine whether related species have similar responses to the same novel pathogen. We found that medium ground finches had a strong transcriptomic response to infection, upregulating genes involved in the innate immune response including interferon production, inflammation, and other immune signaling pathways. In contrast, vegetarian finches had a more limited response to infection. Our results also revealed evidence of viral manipulation of the host’s cellular function and metabolism, providing insight into the ways in which poxviruses affect their hosts. Many of the transcriptomic responses to infection mirrored known processes seen in model and in-vitro studies of poxviruses indicating that many pathways of host defense against poxviruses are conserved among vertebrates and present even in hosts without a long evolutionary history with the virus. At the same time, the variation we observed between closely related species indicates that some endemic species of Galápagos finch may be more susceptible to avian pox than others.

## Introduction

Emerging infectious diseases are a threat to humans and wildlife (Cunningham, Daszak, & Wood, 2017; Daszak, Cunningham, & Hyatt, 2000). Species often vary in their response to novel pathogens, with implications for disease severity, host fitness, and conservation (Zamudio, McDonald, & Belasen, 2020). Understanding the molecular response of hosts to novel pathogens may help explain why disease emerges in some populations and not others, and why some individuals survive while others do not (Liu et al., 2017). The changes to host gene expression following infection are complex and reflect both manipulation of the cellular environment by the pathogen as well as activation of host defense mechanisms (Agudelo-Romero, Carbonell, Perez-Amador, & Elena, 2008; Videvall, Cornwallis, Palinauskas, Valkiūnas, & Hellgren, 2015; Videvall, Palinauskas, Valkiūnas, & Hellgren, 2020). Naïve hosts may have a strong or weak transcriptional response to infection, depending on their ability to detect the pathogen and mount a successful immune response (Videvall et al., 2015). Pathogens also vary in their ability to evade host defenses and replicate in a novel host, depending on how readily they adapt to a new host environment (Agudelo-Romero et al., 2008).

Poxviruses are large, double-stranded DNA viruses of vertebrates and arthropods. Viruses in the genus *Avipoxvirus* infect birds and are distributed worldwide (Williams, Truchado, & Benitez, 2021). Avian pox is transmitted by direct contact between individuals and mechanically by biting arthropods (Zylberberg, Lee, Klasing, & Wikelski, 2012a). Infection typically causes cutaneous lesions on the feet, legs, and face, though in more severe cases it can infect mucous membranes of the respiratory tract (Williams et al., 2021). Mild cutaneous pox infections may not be debilitating; however, large lesions can impede vision, mobility and feeding ability. Avian pox causes more severe disease in insular bird populations that are naïve to the virus (Williams et al., 2021). Pox was implicated in the decline and extinction of several Hawaiian honeycreepers (van Riper III, van Riper, Hansen, & Hackett, 2002)and has recently caused outbreaks of disease in tits (Paridae) in Great Britain (Lawson et al., 2012), magellanic penguins (*Spheniscus magellanicus*) on the Atlantic coast of Argentina (Kane et al., 2012), and endemic passerines (*Calandrella rufescens* and *Anthus berthelotti*) in the Canary Islands (Smits, Tella, Carrete, Serrano, & López, 2005).

Avian pox has been present in the Galápagos Islands for at least a century, but its prevalence may be increasing due to the spread of invasive mosquitos and changes in climate and resource availability (Parker et al., 2011; Zylberberg et al., 2012a; Zylberberg, Lee, Klasing, & Wikelski, 2012b). Pox infects a number of endemic Galápagos passerines including Darwin’s finches (Thraupidae), Galápagos flycatchers (*Myiarchus magnirostris*), and Galápagos mockingbirds (*Mimus spp*.) and has recently been reported in the critically endangered waved albatross (*Phoebastria irrorata*) (Jiménez-Uzcátegui et al., 2019; Tompkins, Anderson, Pabilonia, & Huyvaert, 2017) (Fig 1). Prevalence of avian pox varies among species of Darwin’s finches and may be related to variation in innate immune function (Zylberberg et al., 2012a). Specifically, a previous study that took place between 2000 and 2009 found that prevalence and disease severity was increasing in some populations of ground, tree, and warbler finches (*Geospiza fuliginosa, G. scandens, Camarhychus parvulus* and *Certhida olivacea)*. The increased prevalence of avian pox in these species was correlated with a decrease in PIT54 acute phase protein levels in healthy individuals, suggesting a link between innate immune function and pox susceptibility at the species level. In contrast, medium ground finch (*G. fortis*) populations showed signs of reduced disease spread and increased recovery during the same period and had no change in measures of innate immune function (Zylberberg et al., 2012a).

**Figure 1.**
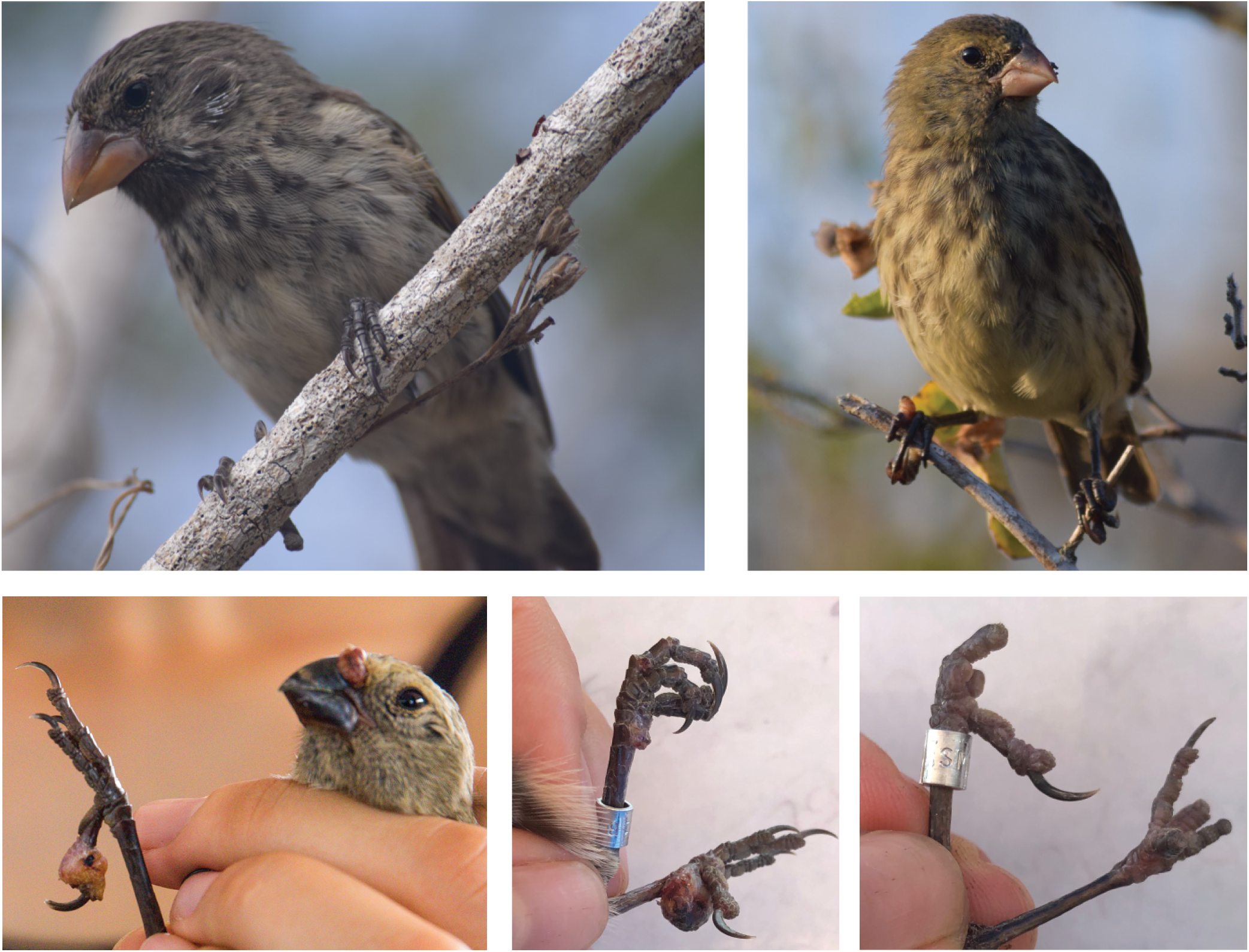
Top row: Medium ground finch (left) and vegetarian finch (right). Bottom row: examples of pox lesions on the feet and nares of a vegetarian finch (left); large lesions on the feet of a medium ground finch (center), and significant digit loss in a ground finch previously infected with pox (right).

Laboratory studies of the model poxvirus vaccinia, the virus used to vaccinate against smallpox, have been instrumental in understanding mechanisms of host defense against poxviruses (Boyle & Traktman, 2009). These studies demonstrate that vertebrate hosts have a complex and robust immune machinery to resist poxviruses. Viral sensing proteins detect viral infection and initiate the innate immune response (Delaloye et al., 2009). Type I interferons are expressed rapidly in response to infection and activate other antiviral responses (Seet et al., 2003). Interferons signal Janus kinases (JAKs) resulting in the phosphorylation of signal transducers and activators of transcription (STATs) that move to the cell nucleus and activate an antiviral state (Seet et al., 2003). The innate immune response also promotes inflammation, recruiting leukocytes to the site of infection (Haga & Bowie, 2005). Cells at the site of infection secrete antiviral and inflammatory cytokines which trigger the adaptive immune response, including the production of antibodies that clear the pathogen and confer lasting immunity (Haga & Bowie, 2005).

In turn, however, poxviruses have sophisticated mechanisms of immune evasion and host manipulation. Poxviruses dedicate a large proportion of their ∼200 genes to encoding immunomodulating proteins (Bidgood & Mercer, 2015). Poxviruses disrupt antiviral defenses by inhibiting the apoptotic response, producing proteins that obstruct interferon and other immune signaling pathways, and co-opting host gene expression (Bidgood & Mercer, 2015; Seet et al., 2003; Smith, Symons, Khanna, Vanderplasschen, & Alcami, 1997; Smith, Talbot-Cooper, & Lu, 2018). In sum, much of the outcome of the pox-host interaction may depend on whether the host or virus is more successful at controlling the intracellular environment and cellular metabolism.

Despite the well-characterized interactions between poxviruses and vertebrate hosts from laboratory experiments, little is known about the transcriptomic effects of pox infection in wild populations. It is unclear whether evolutionary naïve hosts detect pox infection and activate immune responses in an effective way. It is also unknown how effective the poxvirus is at disrupting and evading a new host’s defenses. Mammals appear to have similar immune responses to various poxviruses (Offerman et al., 2015), suggesting that vertebrates could have a conserved reaction to pox infection, even if a particular poxvirus is novel. On the other hand, the fact that avian pox seems to cause particularly severe disease in new host populations suggests that naïve populations of birds may lack effective antiviral defenses, and/or may be particularly vulnerable to viral manipulation.

The goal of this study was to investigate the transcriptomic effects of avian pox infection in two endemic species of Darwin’s finches, medium ground finches *(Geospiza fortis*) and vegetarian finches *(Platyspiza crassirostris*). These focal species are common, sympatric birds in the arid and transitional zones of Santa Cruz Island. We sampled free-living birds and diagnosed finches as actively/currently infected or uninfected based on the presence of distinctive cutaneous pox-like lesions (Parker et al., 2011). Although this method is not a definitive diagnosis, it is a common way of identifying avian pox in the Galápagos (Parker et al., 2011; Tompkins et al., 2017; Vargas, 1987; Zylberberg et al., 2012b). We then used RNA-seq to sequence total mRNA from the blood of individuals with and without pox-like lesions. This technique creates a complete representation of the transcriptome at the time of sampling and thus can identify genes and pathways that are expressed in tandem as well as new candidate genes that may not have had previously known roles in pathogen defense (Videvall et al., 2015).

We used the resulting transcriptomic dataset to address two main questions: 1) whether gene expression differs between finches with and without lesions and 2) whether the response to infection varies between species. To confirm infection with avian poxvirus, we searched the transcriptomic dataset for poxvirus RNA using a metatranscriptomics approach (Galen, Borner, Williamson, Witt, & Perkins, 2020), as well as attempted to amplify pox DNA using PCR. Finally, we tested whether transcriptomic differences birds with and without lesions, and between species, were correlated with immune phenotypes based on leukocyte profiles in the peripheral blood.

## Methods

### Sample collection

Finches were captured by mist-net during January and February of 2019 at the Charles Darwin Research Station on Santa Cruz Island. Birds were visually inspected for the presence of pox lesions and categorized as “infected” if they had visible pox-like lesions on the feet, tarsi, or face (Fig. 1). Lesions were typically accompanied by swelling in the area and occasionally accompanied by necrosis or recent loss of toes. Finches were categorized as “recovered” if they showed signs of previous pox infection (i.e. missing but healed digits). Recovered birds were not included in the genetic study. Finches were scored as “uninfected” if feet and tarsi showed no signs of lesions or swelling and there were no signs of previous pox infection (missing digits). From now on, the terms “infected”, “recovered” and “uninfected” will be used in the paper.

Following capture, birds were banded with a uniquely numbered monel band, and a blood sample (< 75 ul) was taken via brachial venipuncture. A drop of blood was used to make a blood smear for leukocyte profiling. After the smear dried it was fixed in 100% ethanol for one minute and then air dried again. The rest of the blood sample was divided into two equal parts: the first part was immediately preserved in RNA-later. The second part of the sample was preserved on wet ice while in the field. Within 4 hours, the sample preserved on wet ice was centrifuged at 2000 g for 6 minutes to separate the plasma and erythrocytes. The plasma was frozen at −20° C and the erythrocytes were preserved in Queens Lysis Buffer at room temperature. Blood samples preserved in RNA-later were lysed at room temperature for 24 hours, after which they were centrifuged for 10 minutes at 2000 g to compact the blood sample. The supernatant RNA-later was removed with a pipette and the remaining sample was frozen at −20° C. Following the field season, the samples were transported to the Cornell University Museum of Vertebrates where frozen RNA-later and plasma samples were stored at −80° C and lysis buffer samples and blood smears were stored at room temperature.

### RNA extraction

Total mRNA was extracted from RNA-later preserved blood using Qiagen RNeasy kits (Cat #74134) following manufacturer’s protocol. Cells were disrupted by triturating the 20-30 μl of RNA-later preserved blood with the lysis buffer. An additional DNA digestion step was added during extraction using the Qiagen RNase-Free DNase kit following manufacturer’s instructions. RNA extraction quality was verified first using a NanoDrop spectrophotometer to determine concentration and chemical purity (A260/230 and A260/280 ratios) and then on a FragmentAnalyzer (Advanced Analytical) to determine RNA integrity. The RNA quality number (RQN) was > 9.0 for all but one sample, which had an RQN of 7.0.

### Library preparation and sequencing

Forty samples were selected for sequencing: 10 infected and 10 uninfected medium ground finches and 10 infected and 10 uninfected vegetarian finches. Library preparation took place at the Transcriptional Regulation and Expression (TREx) Facility in the Department of Biomedical Sciences, College of Veterinary Medicine, Cornell University. PolyA+ RNA was isolated from total RNA with the NEBNext Poly(A) mRNA Magnetic Isolation Module (New England Biolabs). TruSeq-barcoded RNAseq libraries were generated with the NEBNext Ultra II Directional RNA Library Prep Kit (New England Biolabs). Each library was quantified with a Qubit 2.0 (dsDNA HS kit; Thermo Fisher) and the size distribution was determined with a Fragment Analyzer (Advanced Analytical) prior to pooling. Libraries were sequenced at Novogene on an Illumina NovaSeq 6000 system.

### Bioinformatics

Reads were trimmed for low quality and adaptor sequences with TrimGalore v0.6.0 (https://www.bioinformatics.babraham.ac.uk/projects/trim_galore/), a wrapper for cutadapt (Martin, 2011) and fastQC (https://www.bioinformatics.babraham.ac.uk/projects/fastqc/). We discarded reads shorter than 50 bp and trimmed ends with a quality score < 20, allowing for a maximum error rate of 0.1. The resulting reads were aligned to the *Geospiza fortis* genome archived in NCBI (GeoFor v. 1.0(Li, Li, Parker, & Wang, 2012) using STAR v.2.7.0e (Dobin et al., 2013); reads that did not align to the finch genome were filtered out for metatranscriptomic analysis (below). We used the same reference genome for both species because there is no annotated vegetarian finch genome. We used DESeq2 v.1.26.0 to model normalized counts, calculate log2 fold change between comparison groups, and identify genes that were significantly differentially expressed ((Love, Huber, & Anders, 2014); doi: https://doi.org/10.18129/B9.bioc.DESeq2). The Benjamini–Hochberg method was used to correct p values for multiple testing. Initial PCA analysis of gene expression profiles revealed different profiles for male and female individuals (Supplemental Figure 1). Thus, subsequent Gene Expression Analyses were run on the whole dataset (N = 10 per treatment per species) as well as just for males (ground finch infected N = 5, uninfected N = 7; vegetarian finch infected N = 6, uninfected N = 9). Analyses of the whole dataset excluded sex-linked genes (52 out of 16001 genes).

### Gene Set Enrichment Analysis

We used gene set enrichment analysis (GSEA) to identify biological processes associated with pox infection. GSEA ranks all genes based on their correlation with a relevant phenotype (e.g. infection status) and then tests whether genes in a particular set of interest (e.g. genes involved with a known biological process) are significantly clustered near the top or bottom (Subramanian et al., 2005) We used the log2 fold change estimates generated by DESeq2 to make comparisons between infected and uninfected finches of each species, and between infected vegetarian and ground finches, and uninfected vegetarian and ground finches. We tested for associations between pox phenotype and the 50 Hallmark Gene Sets in the Molecular Signatures Database (Liberzon et al., 2015) and between pox phenotype and the KEGG Orthology Database (M. Kanehisa & Goto, 2000; Minoru Kanehisa, Sato, Kawashima, Furumichi, & Tanabe, 2016). We used the R package clusterProfiler v3.18.1 (Yu, Wang, Han, & He, 2012) to perform gene set enrichment analysis; significant P values were adjusted for multiple comparisons using the “FDR” method. For the Hallmark Gene Sets we used the gene sets collection for chicken (*Gallus gallus*) and for KEGG gene lists we used the collection for medium ground finch. For each database these were the most closely related organisms available for our study species.

### Metatranscriptomics

We screened the transcriptomic dataset for poxvirus reads to verify infection status and quantify viral load. We used Kraken 2 to search for pox reads among all reads that did not align to the ground finch genome (Wood, Lu, & Langmead, 2019). Kraken 2 uses a k-mer-based approach to classify metagenomic sequences and is highly accurate for classifying viruses (Wood et al., 2019). We used the translated search mode and the standard Kraken 2 database which increases sensitivity when searching for viruses and classified sequences at both a confidence level of 0.00 (default) and 0.05 (slightly more stringent. Because poxviruses are diverse and there are no complete avian pox genomes from the Galápagos available, we filtered results for any sequence that was classified to poxvirus subfamily Chordopoxvirinae (poxviruses of vertebrates).

### rtPCR and PCR verification of infection

In an additional attempt to confirm infection status, we used PCR and real-time PCR (rtPCR) to amplify poxvirus DNA. We targeted the *Avipoxvirus* 4b core protein gene following methods described in Baek et al. (2020) (for rtPCR) and MacDonald et al. (2019) (for conventional PCR). DNA was extracted from blood samples preserved at room temperature in Queens Lysis Buffer using Qiagen DNeasy kits. DNA concentration was quantified on a Qubit 2.0 Flourometer (Life Technologies); all samples had a DNA concentration > 5 ng/μL. rtPCR methods followed those described in Baek et al. (2020), with the exception that a standard amount of 25 ng of template DNA was added for all samples. Samples were run in duplicate. Standard PCR methods followed those described in MacDonald et al. (2019); PCR products were then cleaned using a ExoSAP protocol (2 units/μL) (Goldberg & Mason, 2017) and then Sanger sequenced at the Biotechnology Resource Center at Cornell University.

### Leukocyte quantification

We characterized immune phenotypes of finches using slide microscopy of blood smears. We profiled leukocytes in the peripheral blood of 40 ground finches (20 infected and 20 uninfected) and 46 vegetarian finches (24 infected and 22 uninfected). Smears were stained in wright-geimsa stain for 10 minutes (or until cells were clearly visible). Smears were examined by one author (CA), who was blind to species and pox status, under oil immersion at 1000 x magnification until at least 50 leukocytes or 10000 erythrocytes were counted (median leukocytes = 100; median erythrocytes = 11,270). Leukocytes were identified as lymphocytes, monocytes, eosinophils, or heterophils. We calculated the number of each type of leukocyte per 1000 erythrocytes, as well as the heterophil to lymphocyte ratio, a common indicator of stress in birds (Gross & Siegel, 1983). We confirmed that counts were consistent by initially screening a set of ten slides twice, in random order, and estimating repeatability using the lmm method in the R package rptR (v. 0.9.22). Repeatability for lymphocytes, monocytes, and total leukocytes was > 0.80. Basophils, eosinophils, and heterophils were rare or absent from most samples so we did not estimate repeatability for those cell types. We tested for differences in leukocytes per 1000 erythrocytes, lymphocytes per 1000 erythrocytes, monocytes per 1000 erythrocytes, and the heterophil:lymphocyte ratio using generalized linear models with a quasi-Poisson distribution (to account for overdispersion). We originally included sex as a covariate; however, it was not significant in any model so it was removed. We included species and infection status as fixed effects in the final models.

## Results

### RNA-seq results

We sequenced a total of 2.9 billion paired end reads with an average of 74.5 million reads per individual. Across all samples the mean percent alignment to the reference genome was 83.0% (range = 74.8% – 86.5%). There was no significant difference between species in either the raw number or the percent of reads that aligned to the reference genome (linear model P > 0.10 for both raw number and percent alignment).

### Differentially expressed genes

We tested for differential expression of 16,001 genes between various comparison groups: infected vs. uninfected finches for each species, infected ground vs. infected vegetarian finches, and uninfected ground vs. uninfected vegetarian finches. We also repeated the analysis to only include male finches, to eliminate potential confounding effects of sex (Table 1). Between 45 and 1315 genes were significantly differentially expressed between comparison groups (Fig. 2, Table 1). Fewer differentially expressed genes were identified between groups when only males were included (Table 1). More differentially expressed genes were found between infected and uninfected ground finches than between infected and uninfected vegetarian finches.

**Table 1.** Numbers of significantly differentially expressed genes (DEG) between comparison groups

|  | DEG (males and<br>females) | DEG (males only) |
| --- | --- | --- |
| Infected ground vs. uninfected ground | 80 | 35 |
| Infected vegetarian vs. uninfected vegetarian | 45 | 0 |
| Infected ground vs. infected vegetarian | 1157 | 1413 |
| Uninfected ground vs. uninfected vegetarian | 1314 | 969 |

Sample size: N = 10 individuals per group
Sample size males only: ground finch infected N = 5, uninfected N = 7; vegetarian finch infected N = 6, uninfected N = 9.

**Figure 2.**
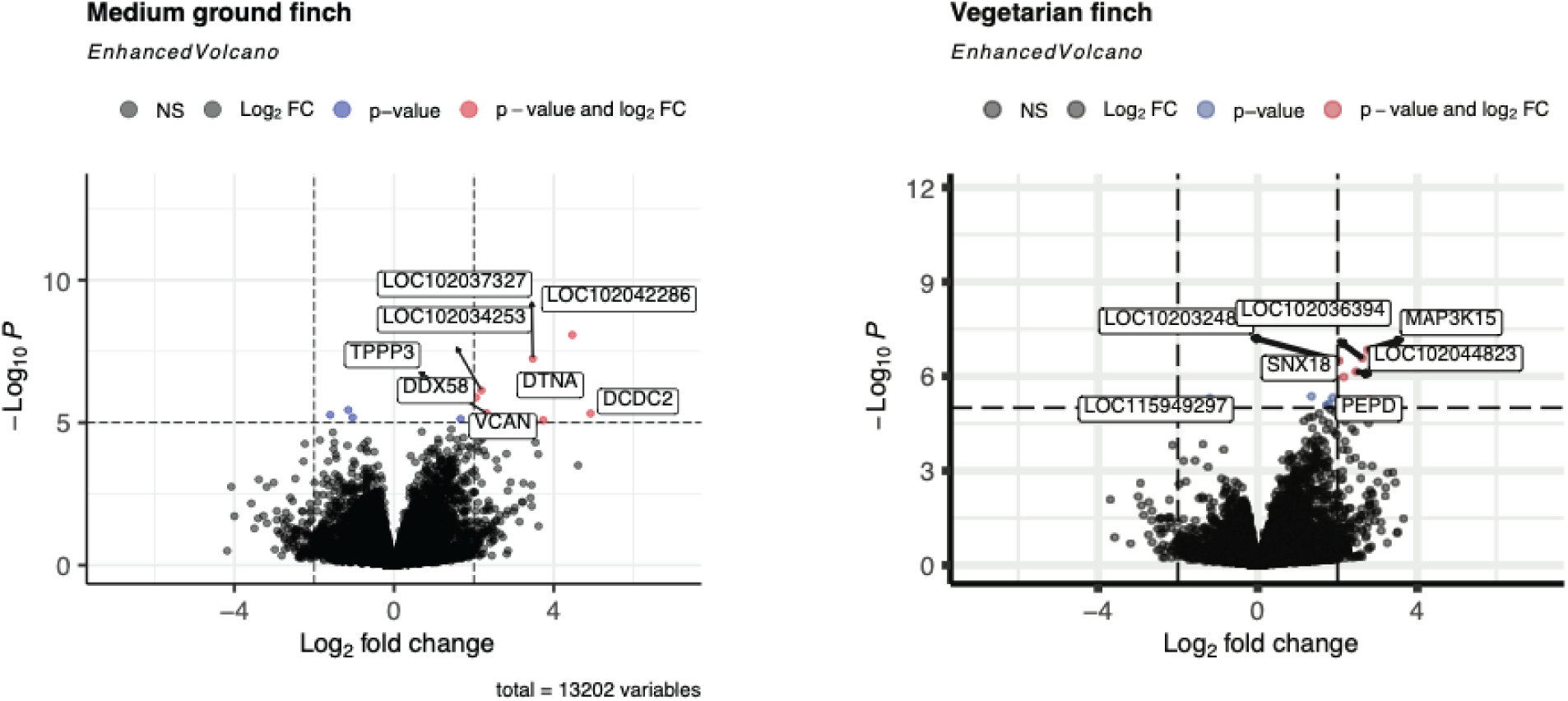
Volcano plot of differentially expressed genes in infected vs. uninfected ground finches (left) and infected vs. uninfected vegetarian finches (right). Each point is a gene. The x axis displays the log_2_fold change in expression in infected birds compared to uninfected birds. The y axis displays the p-value of a Wald test comparing expression of each gene between infected and uninfected groups.

### Gene set enrichment analysis

#### Infected vs. uninfected ground finches

We identified 14 hallmark sets that were significantly enriched in the expression profiles of infected and uninfected ground finches (Fig. 3A). Among these were several sets associated with innate immune function including *interferon alpha response* and *interferon gamma response, IL6 JAK STAT3 signaling, complement, inflammatory response*, and *allograph rejection*. All pathways were upregulated in infected ground finches.

**Figure 3.**
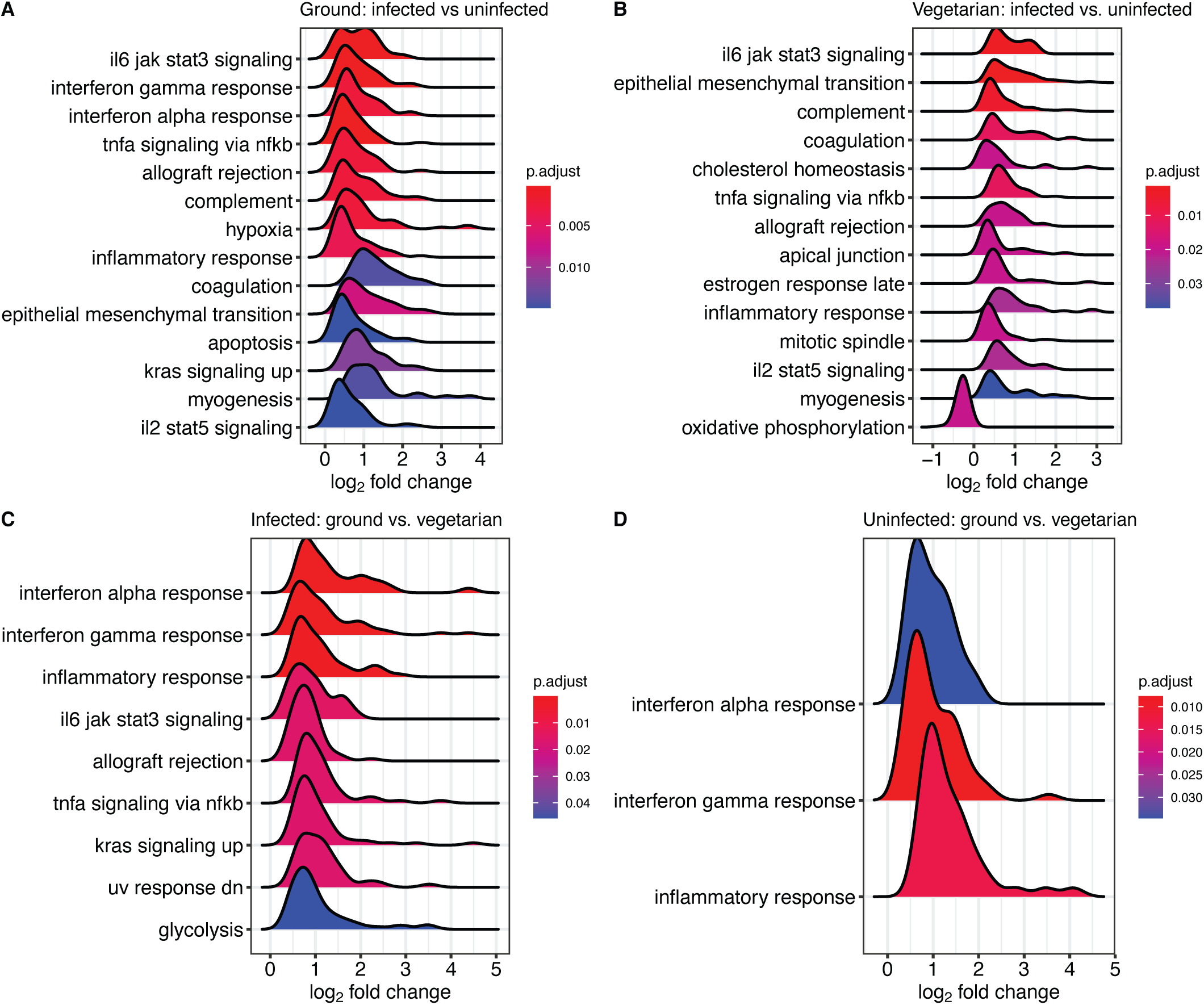
Hallmark gene sets significantly enriched between different comparison groups: A) infected vs. uninfected ground finches, B) infected vs. uninfected vegetarian finches, C) infected ground vs. infected vegetarian, and D) uninfected ground vs. uninfected vegetarian. The x-axis displays the frequency distribution of log_2_ fold changes of genes in that set. The color of each set displays the FDR-adjusted p-value testing whether expression of genes in that set is significantly associated with infection phenotype.

Including only male ground finches, we identified 8 hallmark sets, 7 of which were included in the larger dataset (Fig. S2A). The only set not identified in the whole dataset was *apical surface*, a set related to control of cell polarity in the generation of epithelial cells. The sets with the strongest support between the two analyses were *interferon alpha response, interferon gamma response, allograph rejection*, and *IL6 JAK STAT3 signaling*.

We identified three KEGG pathways that were significantly upregulated in infected ground finches: *Toll-like receptor signaling pathway, Influenza A*, and *phagosome* (Fig. 4A). Two different KEGG pathways were significantly enriched between male infected and uninfected ground finches: *phototransduction* and *ribosome* (Fig. S3A).

**Figure 4.**
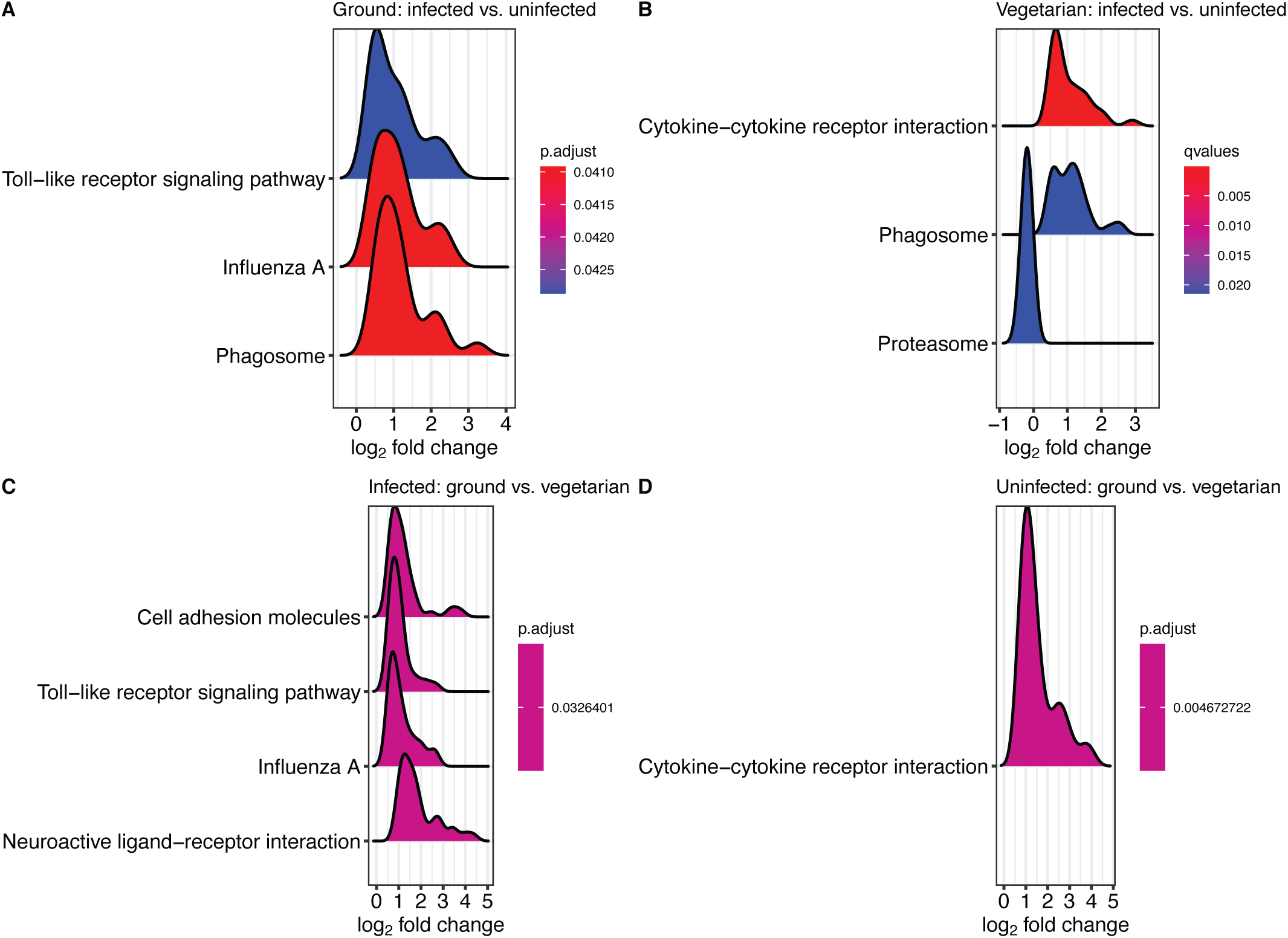
KEGG gene pathways significantly enriched between different comparison groups: A) infected vs. uninfected ground finches, B) infected vs. uninfected vegetarian finches, C) infected ground vs. infected vegetarian, and D) uninfected ground vs. uninfected vegetarian. The x-axis displays the frequency distribution of log_2_ fold changes of genes in that set. The color of each set displays the FDR-adjusted p-value testing whether expression of genes in that set is significantly associated with infection phenotype.

#### Infected vs. uninfected vegetarian finches

Fourteen hallmark sets were significantly enriched between infected and uninfected vegetarian finches (Fig. 3B). Overall, the p-value support was weaker for hallmark set enrichment compared infected vs. uninfected ground finches. Several of the same sets were upregulated in both species in response to infection, including *IL6 JAK STAT3 signaling, epithelial mesenchymal transition*, and *complemen*t. However, other sets were significantly enriched in just vegetarian finches, including *cholesterol homeostasis* and *mitotic spindle*. One set was significantly downregulated in infected vegetarian finches compared to uninfected vegetarian finches: *oxidative phosphorylation*. Limiting the analysis to just males, however, produced zero significant hallmark sets between infected and uninfected groups, potentially because of the reduced sample size.

Three KEGG pathways were significantly enriched between infected and uninfected vegetarian finches: *cytokine-cytokine receptor interaction, phagosome*, which were upregulated in infected finches, and *proteasome*, which was downregulated in infected finches (Fig 4B). However, limiting the analysis to just male finches did not find any significantly enriched KEGG pathways.

#### Infected ground vs. infected vegetarian finches

Compared to infected vegetarian finches, infected ground finches upregulated expression of genes significantly associated with 8 hallmark sets, most notably *interferon response* and *inflammatory response* (Fig. 3C). Similar results were found just comparing infected males of each species (Fig. S2B).

Four KEGG pathways were significantly enriched between infected ground and infected vegetarian finches: *cell adhesion molecules, toll-like receptor signaling pathway, influenza A*, and *neuroactive ligand-receptor interaction* (Fig 4C). Only one KEGG pathway was significantly enriched between infected male ground vs. infected male vegetarian finches: *ribosome*, which was downregulated in ground finches (Fig. S3B).

#### Uninfected ground vs. uninfected vegetarian finches

There were also significant differences in expression between uninfected finches of each species. Three hallmark sets were significantly enriched between ground and vegetarian finches: *interferon alpha, interferon gamma*, and *inflammatory response* (Fig. 3D). Of those three, only *inflammatory response* was significant comparing uninfected males of each species (Fig. S2C).

Only one KEGG pathway was significantly enriched in uninfected ground finches compared to uninfected vegetarian finches: *cytokine-cytokine receptor interaction* (Fig 4D). Analyzing only male finches identified two significant KEGG pathways: *cytokine-cytokine receptor interaction* and *neuroactive ligand-receptor interaction*, both of which were upregulated in ground finches compared to vegetarian finches (Fig. S3C).

### Metatranscriptomics

Very few sequences were classified as poxviruses. The number of reads assigned to the chordopoxvirinae family in each sample ranged from 44 to 514 under the default confidence criterion. Under a slightly higher stringency criterion of 0.05, the number of reads assigned to chordopoxvirus ranged between 0 and 17. There was no significant difference in the number of poxvirus reads either between species or between infected/uninfected birds using either the default or more stringent classification standard.

### rtPCR

No sample was positive for poxvirus using the rtPCR screening method. Most samples (33/40) positively amplified using conventional PCR; however, sequencing results were poor and only two out of 40 samples’ sequences were a match to poxvirus when searched against the NCBI Genbank nucleotide database. These sequences were a 100% match to avipoxvirus 4b core protein sequences from Galápagos, Hawaiian, and North American birds (Gyuranecz et al., 2013). Both samples were from ground finches classified as infected. Other readable sequences were typically short (< 100 bp) and aligned most closely to avian sequences on Genbank.

### Leukocyte profiles

Slide screening recovered between 5 and 247 leukocytes per sample (median = 100). The most common types of leukocytes were monocytes and lymphocytes; eosinophils and basophils were the least common. There were no significant differences in leukocyte counts between infected and uninfected individuals (p > 0.05 for all cell types). Ground finches had significantly more total leukocytes than vegetarian finches (glm p = 0.04; Fig. 5A). On average, ground finches also had more lymphocytes and more monocytes than vegetarian finches; however, the differences were not statistically significant (lymphocytes: glm p = 0.13, Fig. 5B; monocytes: glm p = 0.19, Fig. 5C). There was no significant difference in heterophil:lymphocytes between species (glm p = 0.24, Fig. 3D).

**Figure 5.**
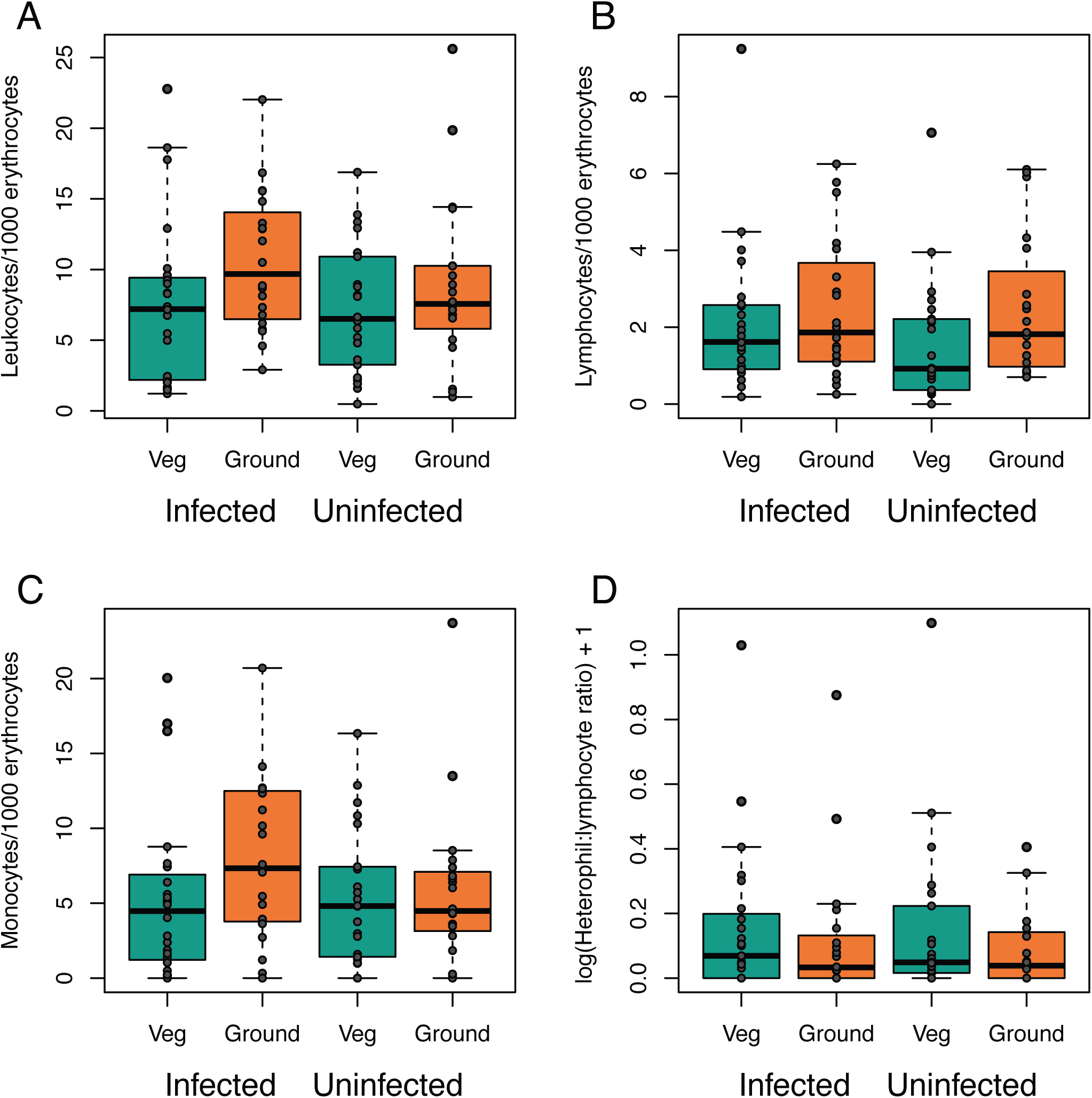
Leukocyte counts of infected and uninfected ground and vegetarian finches: A) total leukocytes, B) lymphocytes, C) monocytes and D) heterophil:lymphocyte ratio. A-C are counts per 1000 erythrocytes.

## Discussion

### Upregulation of genes related to host defense against infection

Infected finches of both species upregulated genes involved in immune function. Infection in ground finches was strongly associated with the upregulation of interferon and interferon-activated genes (Fig 3A). Interferon is a first line of defense against viral invasion (Guerra Susana et al., 2007; Sen, 2001). Type I interferons (including interferon alpha) are released into the intercellular environment in response to viral infection where they initiate a signaling cascade through the JAK-STAT pathway, which activates a network of hundreds of interferon-stimulated genes (Schneider, Chevillotte, & Rice, 2014). The *IL-6 JAK STAT3 signaling* hallmark set was enriched in infected finches of both species, suggesting that the interferon response is disseminated in infected finches. Other KEGG and hallmark sets associated with Type I interferons: *toll-like receptor signaling, TNF-α signaling via NF-κB*, and *apoptosis*, were upregulated in infected ground finches (Fig. 3A, 4A). Toll-like receptors and other viral-sensing proteins detect pathogen infection and trigger the production of interferons (Perdiguero & Esteban, 2009). Tumor necrosis factor (TNF), a cytokine activated by interferon, stimulates the production of nuclear factor kappa B (NF-κB), a transcription factor that is heavily involved in the innate immune response, including by promoting cytokine signaling and apoptosis (Haga & Bowie, 2005). Apoptosis is induced to stop infected cells from producing more virus (Haga & Bowie, 2005; Perdiguero & Esteban, 2009). The hallmark set *interferon gamma* was also enriched in infected ground finches. Interferon gamma is a type II interferon and is produced by natural killer cells and T cells during both the innate and adaptive phases of antiviral response (Sen, 2001). Expression of interferon gamma is promoted by interleukin (IL)-2 (Sen 2001), appears in the upregulated hallmark set IL-2 STAT5 signaling (Fig 3A).

Transcriptomic responses to infection were also associated with generalized immune responses, indicated by the upregulation of genes in the hallmark sets *coagulation, complement, allograft rejection*, and *inflammatory response* and in the KEGG pathways *phagosome* and *cytokine-cytokine receptor interaction* in infected ground and vegetarian finches. Together, these results indicate that Darwin’s finches respond to poxvirus infection using known antiviral pathways. The hallmark and KEGG gene sets that were significantly enriched in infected birds suggest that finches do detect pox infection, respond by producing interferon, a first line antiviral defense, and disseminate the immune response by activating various interferon-stimulated genes.

### Differences between species in immune response to infection

Although both species upregulated genes involved in immune defense, ground finches had a stronger response to infection that vegetarian finches (Fig. 3C). Compared to infected vegetarian finches, infected ground finches had higher expression of genes associated with hallmark sets *interferon alpha* and *gamma* and other immune sets including *inflammatory response, allograph rejection*, and *IL-6 JAK STAT3 signaling*. Our results suggest that poxvirus may be more successful at manipulating or evading defenses of vegetarian finches compared to ground finches. The *toll-like receptor signaling* KEGG pathway was upregulated in infected ground finches; however, it was not significantly enriched in infected vegetarian finches. Thus, one possibility is that vegetarian finches have a more limited ability to detect viral infection compared to ground finches.

Poxviruses have evolved sophisticated mechanisms to evade and block interferon (Bidgood & Mercer, 2015; Seet et al., 2003). Poxvirus can disrupt interferon by disrupting signals from viral-sensing proteins that lead to the transcription of interferon, by competitively binding interferon, disrupting the signaling cascade, and/or by inhibiting the antiviral activity of interferon-stimulated genes (Smith et al., 2018). Fowlpox virus is highly effective at blocking type I interferon responses in poultry and disrupting the interferon signaling cascade (Giotis et al., 2020; Laidlaw et al., 2013). Ground finches do seem to upregulate interferon and interferon-activated genes in response to infection, so avian pox is not completely effective at disrupting host defense. However, the differences we saw in transcriptomic responses of vegetarian finches suggest there may be significant variation among species in this radiation in their ability to detect or mount an immune response against pox. Moreover, the rising prevalence of avian poxvirus in the Galápagos (Zylberberg et al., 2012a). suggests that this virus could be increasingly adapting to the Darwin’s finches as hosts.

Curiously, we found some immune hallmark sets were significantly enriched between uninfected vegetarian and uninfected ground finches. Compared to uninfected vegetarian finches, uninfected ground finches had higher expression of genes involved with interferon alpha, interferon gamma, and the inflammatory response. One potential explanation for this result may be that poxvirus is endemic in our study area and most individuals in the population could have been exposed to the virus. We diagnosed individuals as “infected” or “uninfected” based on the presence of pox-like lesions. However, birds without lesions may have been exposed and/or infected to poxvirus but could have effectively controlled the infection before developing lesions. Indeed, we found that ground finches had significantly higher leukocyte counts than vegetarian finches, regardless of infection status. These data suggest that ground finches could have an earlier and stronger immune response to pox infection before lesions even develop.

### Evidence of viral manipulation of the host cellular environment

Other hallmark and KEGG sets that were significantly enriched in infected birds provide insight into other effects of poxvirus on the host. The hallmark set *epithelial mesenchymal transition* was enriched in both infected ground and infected vegetarian finches compared to uninfected finches (Fig. 3A and B). Epithelial-mesenchymal transition (EMT) is the process through which epithelial cells shift to a phenotype that is non-polarized, migratory, and resistant to apoptosis (Kalluri & Weinberg, 2009). This transition is triggered by signaling pathways including inflammation, hypoxia, KRAS and nuclear factor kappa B (NF-κB) (Lin & Wu, 2020), which were also associated with infection (Fig. 3A and B). EMT is associated both with tissue regeneration as well as oncogenesis (Gonzalez, Costa, Andrade, & Medrado, 2016; Kalluri & Weinberg, 2009). Inflammation triggers EMT, generating fibroblasts, which reconstruct damaged tissues (Kalluri & Weinberg, 2009). Ordinarily, this type of EMT ceases once inflammation reduces (Gonzalez et al., 2016). However, EMT allows dysregulated cells to migrate and invade new tissues, and thus upregulation of EMT is also a hallmark of cancer metastasis (Kalluri & Weinberg, 2009). Several viruses, including herpesviruses, papillomaviruses, and hepatitis viruses promote tumorigenesis by initiating or amplifying EMT (Cyprian, Al-Farsi, Vranic, Akhtar, & Al Moustafa, 2018; Krump & You, 2018). Poxviruses have not historically been included in this group (Cyprian et al., 2018); however, case studies have speculated that avian poxviruses may indeed have oncogenic properties (Pesaro, Biancani, Fabbrizi, & Rossi, 2009; Tsai et al., 1997). The upregulation of genes associated with EMT in infected finches thus may reflect both mechanisms of lesion healing as well as potentially dangerous cancer-related consequences of viral infection.

The upregulation of genes in the cholesterol homeostasis hallmark set in infected vegetarian finches is another potential sign of host manipulation by the virus. Viruses manipulate host membranes and lipid environment both to gain entry to the cell as well as for viral assembly (Deng, Almsherqi, Ng, & Kohlwein, 2010; Heaton & Randall, 2011). Cholesterol is a particularly important lipid for viruses and a lack of cholesterol can inhibit poxvirus replication (Chung, Huang, & Chang, 2005; Deng et al., 2010). Interestingly, the disruption of cholesterol pathways may also interfere with the interferon-regulated JAK-STAT signaling pathway (Mackenzie, Khromykh, & Parton, 2007) reducing the ability of the host’s cells to respond to and signal infection. Infected vegetarian finches did upregulate genes in the Hallmark *IL-6 JAK STAT3 signaling* set; however, fewer immune-related gene sets were significantly upregulated in vegetarian finches compared to ground finches overall. If the poxvirus effectively manipulates cholesterol homeostasis in vegetarian finches, it may be able to concomitantly interfere with the host immune response.

### Cellular hypoxia associated with infection

Infection was associated with changes to the oxygen environment in the host, which may relate to the fight between the host and the virus for control of cellular metabolism (Huang, Huestis, Gan, Ooi, & Ohh, 2021). Infected ground finches upregulated genes involved in hypoxia while infected vegetarian finches downregulated genes involved in oxidative phosphorylation. Hypoxia is a common feature of sites of inflammation because of the increased metabolic demand required to synthesize cytokines, anti-microbial enzymes, and support leukocyte activity (Taylor, Doherty, Fallon, & Cummins, 2016). Hypoxic conditions will trigger a stress response in the cell which is mediated by hypoxia-inducible factor (HIF-1α), a transcriptional regulator of oxygen homeostasis (Rius et al., 2008). Expression of HIF-1α requires NF-κB, a transcription factor that also has an important role in the expression of pro-inflammatory cytokines (Rius et al., 2008). NF-κB signaling was upregulated in infected ground finches, further supporting a link between a hypoxic response and immune activation in infected ground finches.

At the same time, viruses themselves may manipulate the oxygen environment of the host. Viruses activate glycolysis and glutamine metabolism and decrease oxidative phosphorylation to supply energy and biomolecules for viral reproduction (Goodwin, Xu, & Munger, 2015; Thyrsted & Holm, 2021). One way that viruses may force a shift to anerobic metabolism is by inducing a hypoxic response in the host cell. Vaccinia virus interferes with prolyl hydroxylase domain 2 (PHD2), an oxygen sensing enzyme, stabilizing HIF-1α, and leading to a hypoxic response, which may promote viral replication (Huang et al., 2021; Mazzon et al., 2013). Although the hypoxia hallmark set was not significantly enriched in infected vegetarian finches compared to uninfected vegetarian finches, infected vegetarian finches downregulated genes involved in oxidative phosphorylation, suggesting that the virus may be manipulating cellular metabolism towards anaerobic metabolism. Upregulation of genes in the hallmark set *TNF-α signaling via NF-κB* was stronger in infected ground finches compared to infected vegetarian finches (Fig. 3A, B). Since the cellular response to hypoxia also depends on the inflammatory cytokine NF-κB, a weaker cytokine response could explain why we did not observe a similar change in hypoxia genes in vegetarian finches.

### Differences in gene expression between sexes

We chose to sequence infected birds with the most visible pox infections regardless of sex, so our dataset included both males and females. A PCA of overall gene expression revealed differences between species but also between male and female birds (Fig. S1). Many of these differences likely resulted from the fact that we sampled birds during the breeding season during which the birds undergo sex-specific hormonal changes. Males and females were distributed between infected and uninfected groups (Fig. S1) and when we tested for differences in expression between infected and uninfected finches of both sexes, most of the gene sets that were significantly enriched had a clear relationship to antiviral immune function. Thus, we have no reason to suspect that our results were confounded with sex. Still, we repeated our analysis including just male birds (which comprised most of the samples). The results we found were broadly similar for ground finches (Fig. S2, S3). However, we did not find any significant hallmark sets between infected and uninfected vegetarian finch males (Fig. S2). This result was surprising since only 5/20 samples were from female birds.

It is possible that pox infection affects males and females differently, especially during the breeding season. Males and females both provision nestlings but also engage in distinct reproductive activities: males invest energy in territory defense and nest construction, females produce and incubate eggs. Male animals tend to be more susceptible than females to parasites and pathogens, in part due to the effects of androgens on the immune system (Klein, 2000). Thus, we could expect sex-based differences in the transcriptomic response to pox. However, we did not have enough power to test for an interaction between transcriptomic response to infection and sex. Nevertheless, our results analyzing male finches alone are consistent with the conclusion that ground finches upregulate innate immune genes in response to pox infection and that vegetarian finches have a more limited transcriptomic response to pox.

### Study limitations

We observed significant differences between species in their response to pox infection. However, experimental infections are not permissible in this system, so we had to rely on sampling birds captured with pox-like lesions. Correspondingly, our results are correlative and likely reflect a general response of finches after several days of infection. The transcriptomic responses of the finches to poxvirus may be different during the acute phase of infection (i.e. in the first hours following infection).

It is possible that the differences we observed between species were artifacts of natural variation in infection intensity. For instance, if ground finches were infected more recently than vegetarian finches, or if we happened to sample ground finches with more intense infections, we could have observed a stronger transcriptomic response in ground finches by chance. However, our two study species are syntopic, forage in similar areas, and were frequently captured at the same time, thus, we have no reason to suspect differences in pox exposure. Experimental infections in a captive, comparative system would provide additional insight into the dynamics of the host transcriptomic response to avian pox infection. Nevertheless, our results provide a first glimpse into the transcriptomic effects of this emerging disease in free-living birds.

The presence or absence of lesions is not a definitive diagnosis of avian pox infection. We attempted to confirm infection and quantify viral load through several different methods: metatranscriptomics, rtPCR, and conventional PCR. However, none of these methods successfully amplified poxviral RNA or DNA from the blood samples of birds with lesions. We do not take these data to mean that our birds were uninfected with pox. Diagnosis of poxvirus infection or the absence thereof remains challenging in wild birds (Farias et al. 2010; Baek et al. 2020). Poxviruses tend not to replicate in blood cells, so definitive diagnosis often requires isolating live virus and/or using histopathology to confirm the presence of viral inclusion bodies (Bollinger bodies). However, biopsy of lesions from free-living birds is not always advisable because it creates an open wound and presents opportunities for secondary infection (Farias et al. 2010) and histopathology cannot be used to determine if a bird is uninfected. PCR and real-time PCR have emerged as possible ways to screen birds for poxvirus infection; however, PCR results do not always agree with histopathology diagnoses or presence of lesions (Baek et al., 2020; Farias et al., 2010; Parker et al., 2011). For these reasons, the presence/absence of pox-like lesions is frequently used to diagnose avian pox in the Galápagos since there are no other documented diseases present that would cause similar pathologies (Kleindorfer & Dudaniec, 2006; Lindstrom, Foufopoulos, Parn, & Wikelski, 2004; Parker et al., 2011; Zylberberg et al., 2012b, 2012a).

We had hoped that screening our RNA-seq dataset for viral reads would reveal useful information on viral prevalence and intensity since metatranscriptomics approaches have produced genomic data from other parasites and pathogens (Galen et al., 2020; Shakya, Lo, & Chain, 2019). However, we detected few poxvirus reads, potentially because poxvirus is not known to replicate in blood cells. Even though we did not detect poxvirus in the blood, we did observe transcriptomic differences in the blood between birds with and without pox lesions. These transcriptomic changes may have occurred in leukocytes as they were activated by infection and recruited from general circulation to the site of infection (Chen et al., 2017). It is possible that the transcriptomic effects of pox in infected tissues may be different, and more reflective of viral manipulation of the host’s cellular machinery. Still, our data indicate that some effects of poxvirus infection may be observed in non-infected tissues, and that blood may be a useful proxy in cases where wild animals cannot be sampled destructively.

Transcriptomic studies in free-living organisms are important because they can provide novel information about the function of genes that have no analogue in related model organisms (Alvarez, Schrey, & Richards, 2015). Indeed, several of the genes most differentially expressed in response to pox infection were uncharacterized (Fig. 2), meaning we do not have information about their potential function. These genes provide important candidates for further study of host-virus interactions.

### Implications for Darwin’s finch conservation

Sixteen of the 28 native species of land birds in the Galápagos Islands are considered threatened by IUCN (BirdLife International 2020). Invasive parasites and pathogens are important threats to island communities including Galápagos birds. However, the effects of pathogens on a population level can be difficult to quantify without longitudinal data and the ability to accurately identify disease-caused mortality (Cunningham et al., 2017). Indeed, the role of disease in wildlife declines may well be underestimated due to the cryptic nature of the effects of disease (Cunningham et al., 2017). Thus, understanding the mechanisms of susceptibility or resistance to emerging pathogens in Darwin’s finches is important for gauging the threat that parasites and pathogens pose to different populations and allocating resources for conservation and management appropriately. The effects of pathogens are particularly challenging to study in places like the Galápagos where endemic species are highly protected and research-based interventions are restricted. Correspondingly, we had to rely on an observational approach sampling naturally infected birds.

Prevalence of pox lesions varies among species of Darwin’s finches and was previously associated with variation in innate immune function (Zylberberg et al., 2012a). However, this previous work largely focused on ground finches and tree finches, and vegetarian finches were not sampled. Vegetarian finches are from a different clade in the Galápagos finch radiation (Lamichhaney et al., 2015) and they occur in lower density than *Geospiza* ground finches (Dvorak, Fessl, Nemeth, Kleindorfer, & Tebbich, 2012). Relatively little is known about the threats of introduced species to vegetarian finch populations (Heimpel, Hillstrom, Freund, Knutie, & Clayton, 2017). Population censuses completed between 1997 and 2010 found that vegetarian finch populations were stable in the arid zone of Santa Cruz (where our sampling took place) and declining in the humid agricultural zone (Dvorak et al., 2012). In contrast, medium ground finch populations increased in the arid zone and were stable in the agricultural zone. Over the same period, the prevalence of avian pox increased in the agricultural zone (Zylberberg et al., 2012b). Neither of our study species is very abundant in the agricultural zone. However, it is possible that avian pox is contributing to differences in population trends between these two species. However, future studies using mark-recapture methods could test whether pox has differential effects on survival of various species of Galápagos finches.

Avian pox contributed to the decline and extinction of bird populations in Hawaii and has been associated with disease in other novel hosts (van Riper III et al., 2002; Williams et al., 2021). In the Galápagos avian pox is prevalent and a cause for concern; however, little is known about the effects of pox infection on Galápagos bird physiology and cellular function. The results here indicate that wild naïve passerines have similar antiviral transcriptomic responses to poxvirus infection as do mouse and human models. These data are evidence for a conserved response to poxviruses in vertebrates, even in hosts without a long evolutionary history with pox. At the same time, the variation we observed between related species shows that not all naïve hosts respond to a novel infection in the same way and that some Galápagos species may be more vulnerable to pox than others.

## Acknowledgements

The work was done with permission from the Galápagos National Park (PC-30-19) and the Ecuadorian Ministry of the Environment (MAE-DNB-CM-2016-0043). We thank Kiyoko Gotanda, Ashley Saulsberry, and Sarah Wagner for assistance in the field; the Charles Darwin Research Station and Goberth Cabrera for logistical support; Charles Dardia for permit support; Bronwyn Butcher, Jen Grenier, Ann Tate, Faraz Ahmed and the Cornell Transcriptional Regulation & Expression Facility for assistance with library preparation, sequencing, and data processing. This publication is contribution number 2424 of the Charles Darwin Foundation for the Galapagos Islands. This work was funded by a Rose Postdoctoral Fellowship to SMM.

---

**Figure S1.**
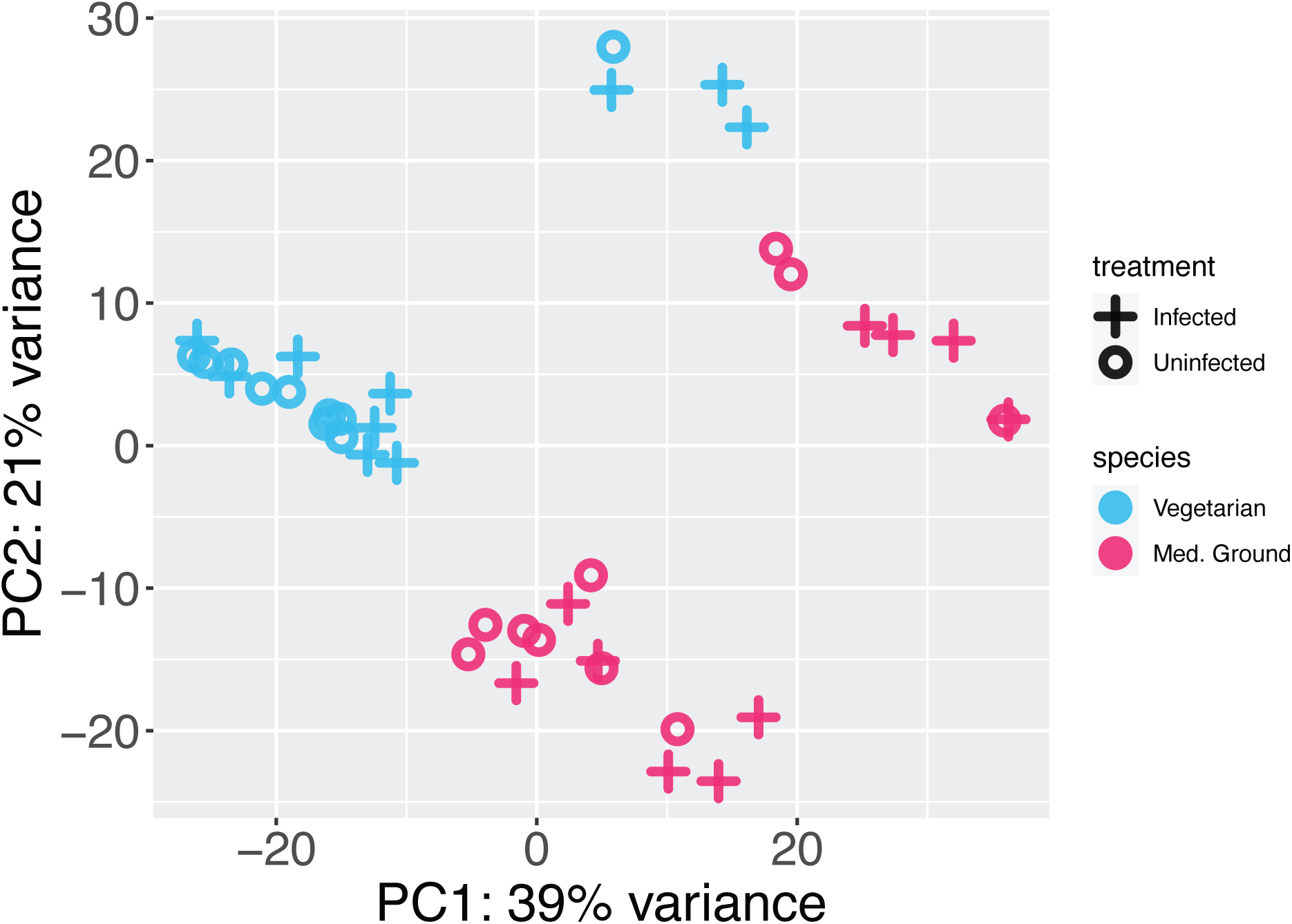
PCA of expression values for individuals in the study. Cluster of individuals in the upper right hand quandrant corresponds to female birds.

**Figure S2.**
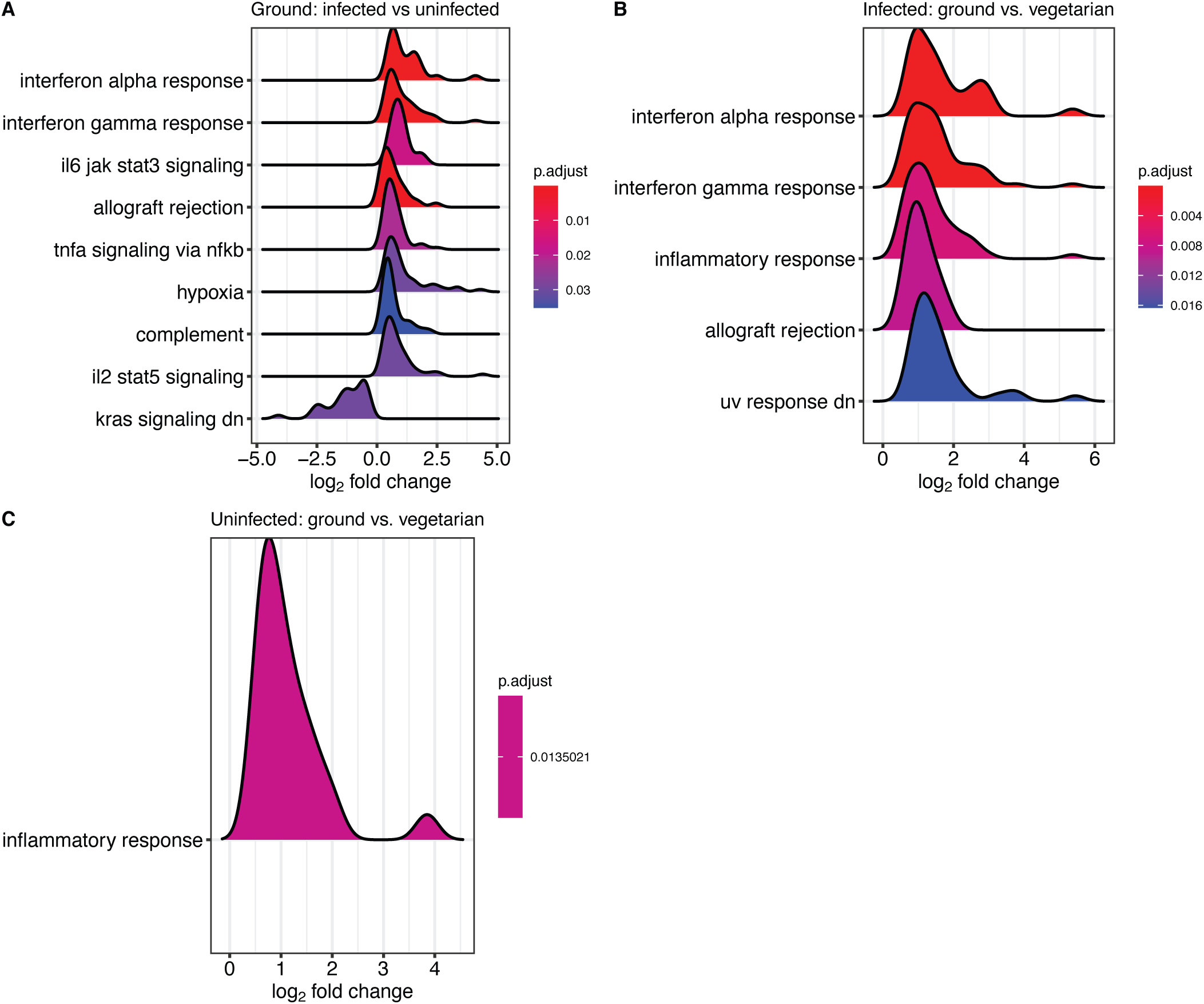
Hallmark gene sets significantly enriched between different comparison groups including only male birds. No gene sets were significantly enriched between infected and uninfected male vegetarian finches.

**Figure S3.**
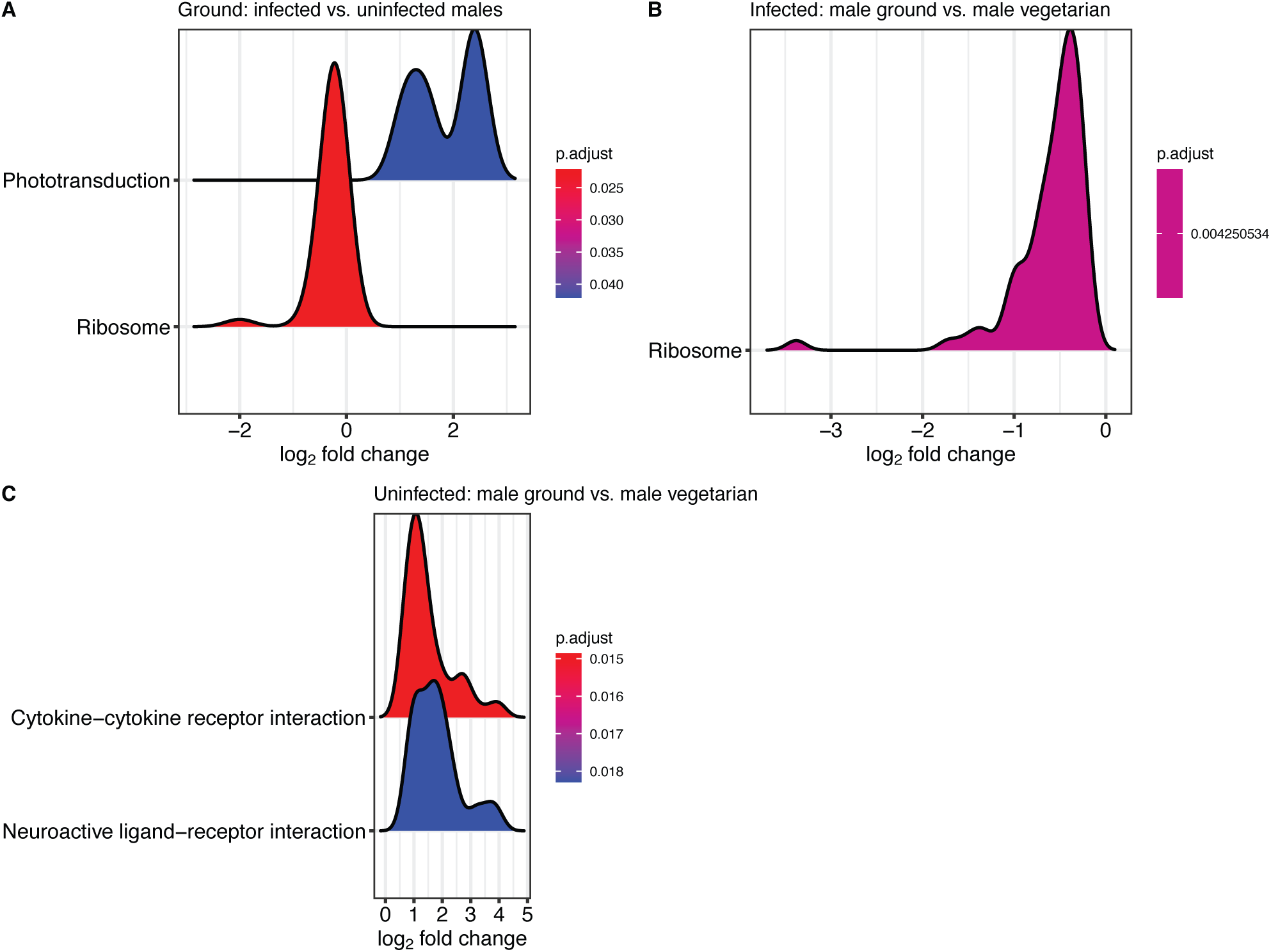
KEGG gene pathways significantly enriched between different comparison groups including only male birds. No gene pathways were significantly enriched between infected and uninfected male vegetarian finches.

## Notes

### Competing Interest Statement

The authors have declared no competing interest.

